# Epileptic encephalopathy-related Kv2.1 mutants impair channel clustering and membrane distribution but not neuronal excitability

**DOI:** 10.1101/2025.10.13.681825

**Authors:** Anne-Lise Paupiah, Melvyn Ginisty, Marion Russeau, Imane Mouktine, Sabine Levi, Jean-Christophe Poncer, Marianne Renner

**Author notes:** Institut du Cerveau, INSERM-Sorbonne Université UMR-S 1127, CNRS U7225, 47 Bvd. De l’Hôpital, 75013 Paris, France. Correspondence to : Marianne Renner, Institut du Cerveau, 47 Bvd. De l’Hôpital, 75013 Paris, France.

## Abstract

The voltage-gated potassium channel Kv2.1, which is encoded by the epileptic encephalopathy-associated gene *KCNB1*, is a primary driver of delayed-rectifier K^+^ currents in neurons. These currents contribute to high frequency firing by preventing depolarization block due to Na^+^ channel inactivation. Wild-type (WT) channels are localized at the soma, proximal dendrites and axons, forming large aggregates (clusters) via their C-terminal proximal restriction and clustering domain (PRC). This study investigates the biophysical and functional consequences of two C-terminal truncation mutations (Y529* and R579*), identified in patients with epileptic encephalopathy, which disrupt this critical clustering domain. Cluster formation was impaired, though not abolished, in neurons expressing mutated subunits together with endogenous WT subunits.

Consistent with this clustering deficit, single-molecule imaging revealed altered channel dynamics, with WT channels remaining largely immobile, while mutated forms exhibited intermittent diffusion punctuated by transient immobilization events. This behavior was reproduced by diffusion-capture simulations considering channels with different number of PRC domains (i.e. with mutant subunits) and labile scaffolding interactions. Patch-clamp recordings revealed no significant difference in excitability between WT and mutant-expressing neurons. However, we observed increased firing in both conditions compared to non-transfected neurons. This suggests that Kv2.1 overexpression (for both WT and mutant channels) counterintuitively enhances neuronal excitability. Our data challenge a canonical role of this voltage-gated potassium channel in dampening neuronal firing, indicating that its overall expression level can be a critical and paradoxical driver of hyperexcitability.

## Introduction

Voltage-dependent potassium (Kv) channels are fundamental to the intrinsic excitability of neurons. Kv2.1 channels are the primary contributors to delayed rectifier (DR) K^+^ currents and are widely expressed in the brain, where they play an important role in neuronal excitability (Murakoshi & Trimmer, 1999; Du et al., 2000; Vacher et al., 2008; Liu & Bean, 2014). These channels have relatively slow activation kinetics that modulate action potential (AP) width and afterhyperpolarization (AHP; Guan et al., 2007; Liu & Bean, 2014). Their primary role is to support high-frequency firing in pyramidal neurons during sustained activity by preventing depolarization block in CA1 neurons (Guan et al., 2013; Liu & Bean, 2014; Steinert et al., 2011). Kv2.1 channels are composed of four α-subunit polypeptides, each with six transmembrane segments (S1–S6).

These channels can form homo-or hetero-tetramers. For instance, they form 2:2 complexes with silent Kv6 (Bocksteins et al., 2016; Möller et al., 2020) and Kv5.1 subunits (Ferns et al., 2025). Within individual neurons, Kv2.1 channels form large aggregates (clusters) in the soma, proximal dendrites, and the axonal initial segment (Sarmiere et al., 2008; Trimmer, 2015). Clustering is mediated by a sequence in the C-terminus, the proximal restriction and clustering domain (PRC; Lim et al., 2000).

The interest of clustering for Kv2.1 function is twofold. First, it has been proposed that Kv2.1 conductance could be downregulated by local channel density, placing clustered channels in a state of low conductance (Fox et al., 2013). Second, several studies have demonstrated that Kv2.1 plays a structural role in clusters; at least some of these clusters co-localize with areas of contact between the endoplasmic reticulum (ER) and the plasma membrane (PM). In these structures, ER-associated VAMP-binding proteins (VAPs) are likely responsible for the scaffolding interactions that immobilize Kv2.1 in clusters (Fox et al., 2015; Kirmiz et al., 2018; Johnson et al., 2019).

A growing number of *de novo* mutations in the Kv2.1 gene, *KCNB1*, have been reported in patients with epileptic encephalopathy (Bar et al., 2020). The effects of these mutations on Kv2 currents have mainly been characterized in heterologous systems, and only a few of their consequences on neuronal excitability are known (Torkamani et al., 2014; Kang et al., 2019). Interestingly, mutations targeting the C-terminal domain lead to truncated proteins lacking the PRC domain (Jensen et al., 2017; Lim et al., 2000; Mohapatra & Trimmer, 2006; Scannevin et al., 1996). Patients with these mutations experience significant cognitive impairment and autism spectrum disorders, though seizures tend to be mild or infrequent (Bar et al., 2020; de Kovel et al., 2017).

We analyzed the consequences of expressing the C-terminal truncation mutants Y529* and R579* of Kv2.1 in cultured hippocampal neurons. Our focus was on the subcellular distribution of the channel and changes in neuronal excitability.

Using single-molecule localization microscopy and single-particle tracking (SPT), we examined the distribution and lateral diffusion of Kv2.1 channels with wild-type (WT) or mutated subunits. Both mutations altered cluster formation without preventing it. SPT experiments revealed that WT channels are essentially immobile, whereas channels with mutated subunits alternate between immobilization in clusters and free diffusion as non-clustered channels. Simulations of diffusion and trapping of molecules with varying numbers of PRC domains reproduced these results, suggesting labile interactions with scaffolding elements and indicating that the formation of large clusters requires channels with two or more subunits carrying PRC domains. Neuronal excitability was not significantly affected by overexpressing mutant channels. However, we observed an unexpected increase in excitability in neurons overexpressing WT or mutant channels, suggesting that changes in channel expression levels, rather than clustering capacity, may control neuronal excitability.

## Materials and methods

### Neuronal culture

All animal procedures were carried out according to the European Community Council directive of November 24^th^ 1986 (86/609/EEC), the guidelines of the French Ministry of Agriculture and the Direction Départementale des Services Vétérinaires de Paris (Ecole Normale Supérieure, Animalerie des Rongeurs, license B 75-05-20). Approval was obtained by the Comité d’Ethique pour l’Expérimentation Animale Charles Darwin (licence Ce5/2012/018). All efforts were made to minimize animal suffering and to reduce the number of animals used. Primary cultures of hippocampal neurons were prepared as previously described (Chamma et al., 2013) with some modifications of the protocol. Briefly, hippocampi were dissected from embryonic day 18 or 19 Sprague-Dawley rats of either sex. Tissue was then trypsinized (0.25% v/v; ThermoFisher Scientific, France), and mechanically dissociated in 1X HBSS (1X; ThermoFisher Scientific, France) containing HEPES (10mM; Invitrogen). Neurons were plated at a density of 180 × 10^3^ cells/ml onto 18-mm diameter glass coverslips (Assistent; Germany) pre-coated with poly-D,L-ornithine (50 µg/ml; Sigma-Aldrich; Lyon, France) in plating medium composed of Minimum Essential Medium (MEM; Sigma-Aldrich) supplemented with horse serum (10% v/v), L-glutamine (2 mM) and Na^+^ pyruvate (1 mM; ThermoFisher Scientific, France). After attachment for 3-4 hours, cells were incubated in culture medium that consists of Neurobasal medium supplemented with B27 (1X), L-glutamine (2 mM), and antibiotics (penicillin 200 units/ml, streptomycin, 200 µg/ml) (ThermoFisher Scientific, France) for up to 4 weeks at 37°C in a 5% CO_2_ humidified incubator. Each week, one third of the culture medium volume was renewed.

### Neuronal transfection and DNA constructs

Transfections were carried out at DIV 9-10 using TransFectin (BioRad; France) according to the manufacturer’s instructions (DNA:TransFectin ratio 1 µg:3 µl), with 1-1.2 µg of plasmid DNA per 20 mm well. Kv2.1WT-GFP and Kv2.1WT-Dendra2 were derived from rKv2.1WT-GFPHis, kindly provided by J. R. Martens (University of Florida, USA). Kv2.1Y529*-GFP, Kv2.1R579*-GFP, Kv2.1Y529*-tomato and Kv2.1R579*-tomato were derived from Kv2.1WT-GFP and Kv2.1WT-Dendra2 by site-directed mutagenesis. Homer-DsRed was a kind gift of D. Choquet and Glycine receptor chimera a1m was kindly provided by A. Triller. Experiments were performed between 21-27 DIV.

### Immunocytochemistry

For detection of surface Kv2.1WT-GFP or GFP-tagged mutants, live neurons were incubated for 10 min at 37°C with alpaca nanobodies against GFP coupled to AlexaFluor647 (GFP-Booster, 1/200; Proteintech; Germany) diluted in the culture medium, followed by two washes in PBS 1X. Neurons were then fixed for 10 min at −20°C in 100% methanol and washed in PBS 1X.

For intracellular labeling, cells were fixed for 15 min at room temperature (RT) in paraformaldehyde (PFA, 4% w/v; Sigma-Aldrich) and sucrose (4% w/v; Sigma-Aldrich) solution prepared in PBS. Following washes in PBS, the cells were permeabilized with Triton (0.25% v/v; Sigma-Aldrich) diluted in PBS. Cells were washed again in PBS and incubated for 30 min at RT in blocking solution (PBS enriched with goat serum, 10% v/v; ThermoFisher Scientific, France). Subsequently, neurons were incubated for 1h with rabbit anti-Kv2.1 antibody (1:100; Alomone Labs, Israel) diluted in blocking solution and 45 min at RT with the corresponding secondary antibody (AlexaFluor647-or Cy3-tagged donkey anti-rabbit; Jackson Immunoresearch, USA). After final rinsing, coverslips were mounted on glass slides using Mowiol 4-88 (48 mg/ml; Sigma-Aldrich). Sets of neurons compared for quantification were labeled simultaneously.

### Fluorescence image acquisition and analysis

Confocal imaging was performed on a Leica TCS SP5 (Leica Microsystems, Germany) equipped with a 100X oil-immersion objective, acquiring optical sections of 0.25 µm in depth.

For wide-field imaging, image acquisition was performed using a 63X oil-immersion objective (NA 1.32) on a Leica DM6000 upright epifluorescence microscope (Leica Microsystems) with a 12-bit cooled CCD camera (Micromax, Roper Scientific, France) run by MetaMorph software (Roper Scientific). Quantification was performed using custom software written in MATLAB (The MathWorks, USA). Camera exposure time was determined on bright cells to obtain the best fluorescence-to-noise ratio and to avoid pixel saturation. All images from a given culture were then acquired with the same exposure time and acquisition parameters.

Surface staining was quantified on maximal projections (confocal images, typically 2-3 optical slices) or fluorescence images (widefield) in which three regions of interest (ROI) were manually chosen at different locations on a dendrite. Proximal, intermediate, and distal ROIs were placed at ~50, ~100 and ~150 µm from the soma, respectively, using tools provided by Fiji Image analysis software. Average fluorescence was quantified in each ROI divided by the mean of proximal ROIs for normalization.

For clustering analysis on confocal images, background was flattened using the “Subtract Background” tool in Fiji (https://fiji.sc/). Residual background was subtracted from the whole image before applying an intensity threshold for segmentation. Segmented images were used as a mask to recover total fluorescence values in the original image and to extract the area of clusters, using home-made custom routines in MATLAB.

### Super-resolution imaging

Super-resolution imaging on fixed samples was conducted on an inverted N-STORM Nikon Eclipse Ti microscope (Nikon Instruments, Netherlands) with a 100X oil-immersion objective (NA 1.49) and an Andor iXon Ultra 897 EMCCD camera (image pixel size, 105 nm, Ireland), using specific lasers for STORM imaging of AlexaFluor647 (405 and 640 nm). Samples were imaged in a PBS-based oxygen-scavenging medium containing imaging buffer (Tris 100 mM, NaCl 20 mM, pH 8), glucose (40% w/v; Sigma-Aldrich), PBS 1X, cysteamine hydrochloride (MEA 100mM; Sigma-Aldrich), catalase (5 mg/ml; Sigma-Aldrich) and pyranose oxidase (200 U/ml; Sigma-Aldrich). Catalase was diluted in MgCl (4 mM), EGTA (2 mM) and PIPES (24 mM; Sigma-Aldrich, pH 6.8). Pyranose oxidase was diluted in the same buffer supplemented with glycerol (50% v/v). Videos of 30 000 frames were acquired at a frame rate of 20 ms. The z position was retained during the acquisition by a Nikon Perfect Focus System.

### Single-molecule localization and clustering analysis

Single-molecule localization and 2D image reconstruction were performed as described in Specht et al., 2013, using a homemade software in MATLAB (SuperRes), available at github.com/mlrennerfr/SuperRes. Briefly, signals from spatially separated fluorophores were adjusted to a 2D Gaussian distribution representing the point spread function of the set-up. Poorly localized peaks (fitting R^2^ < 0.7) were not included for further analysis. The drift of the stage was corrected using 100 nm multicolor fluorescent beads (TetraSpeck, 1/300; ThermoFisher Scientific, France) to follow the movement through the frames; and was nullified by subtracting the displacement of beads from detection coordinates of each frame. Rendered images were obtained by superimposing the coordinates of single-molecule detections, which were represented with 2D Gaussian curves of unitary intensity and SDs representing the localization precision (Baddeley et al., 2010).

Clustering analyses were performed using *Diinamic* (github.com/mlrennerfr/Diinamic), a custom software developed in the lab (Paupiah et al., 2023). Clustering detection was carried out on regions of interest (ROIs) drawn on top of pointillistic images constructed from the coordinates of detections. Detections were initially sorted into belonging to candidate clusters or not depending on their localization with respect to a segmentation mask. This mask was created by thresholding the pixel intensity of the corresponding rendered image. The clusters’ borders were defined by the *boundary* function of MATLAB. Candidate clusters were retained if: 1) their density, calculated in a pixel-wise manner, exceeded a threshold; 2) their sizes were between given thresholds (minimum and maximum). Thresholds were chosen considering the expected number of detections per molecule, the size of the molecule and the expected size of clusters given other microscopy data already published. Briefly, we applied Diinamic-R (intensity threshold 15%) with a minimum density of 2 detections per SMLM pixel and at least 150 detections per cluster.

### Live cell staining for single particle imaging

To track membrane GFP-tagged channels, quantum dots (QDs) were pre-coupled to anti-GFP primary antibodies (rabbit; Roche, Switzerland) as reported previously (Renner et al., 2017). a1m was tracked by QD coupled to a monoclonal anti-myc antibody (Roche). Briefly, goat anti-rabbit F(ab’)2-tagged QDs emitting at 655 nm (Q11422MP; Invitrogen) were incubated first with the antibody for 30 min in PBS, and then blocked for 15 min with casein in a final volume of 10 μL. Neurons were incubated with the pre-coupled QDs (1:6000–1:10000 final QD dilution) for 5 min at 37°C.

### Single particle tracking and analysis

Cells were imaged using an Olympus IX71 inverted microscope equipped with a 60X oil-immersion objective (NA 1.42; Olympus, Japan) and a X-Cite 120Q lamp (Lumen Dynamics, USA). Individual images of Kv2.1-GFP, Homer-DsRed and QD real time recordings (integration time of 75 ms over 1200 consecutive frames) were acquired with Hamamatsu ImagEM EMCCD camera (Hamamatsu Photonics, Japan) and MetaView software (Meta Imaging 7.7; MetaView software).

Tracking was performed with homemade software in MATLAB (SPTrack_v4), available at github.com/mlrennerfr/SPTrack. The center of the spot fluorescence was determined by a 2D-Gaussian fit. Spatial resolution was ~10-20 nm. The spots in a given frame (time point) were associated with the maximum likelyhood trajectories estimated on previous frames of the image sequence. In some experiments, trajectories were defined as belonging to a cluster if they were on top of clusters defined by a binary mask of Kv2.1WT-GFP fluorescence. The mean square displacement (*MSD*) was calculated using:

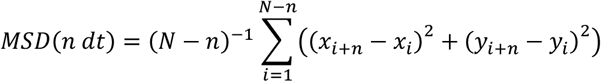

where *x*_*i*_ and *y*_*i*_ are the coordinates of an object on frame *i, N* is the total number of steps in the trajectory, *dt* is the time interval between two successive frames, *n* is the number of frames and *ndt* is the time interval over which displacement is averaged. The diffusion coefficient *D* was calculated by fitting the first 2 to 5 points of the *MSD* plot versus time with the equation:

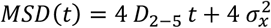

with *σ*_*x*_ the spot localization accuracy (positional accuracy) in one direction.

The packing coefficient (*Pc*) at each time point *i* was calculated as

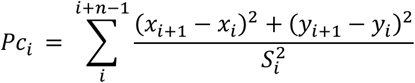

where *x*_*i*_, *y*_*i*_ are the coordinates at time *i*; *x*_*i+1*_, *y*_*i+1*_ are the coordinates at time *i+1, n* is the length of the time window (*n* = 30 time points) and *S*_*i*_ is the surface area of the convex hull of the trajectory segment between time points *i* and *i+n. S*_*i*_ was calculated using the *convhull* function in Matlab (Renner et al., 2017). Immobile molecules display a trajectory that is confined in an area corresponding to the localization precision, thus stabilization events were detected using two criteria: the value of *Pc* corresponding to such confinement, and the size of the confinement area during the event. Given the localization precision in our experiments, we set the threshold to detect stabilization at 4500 µm^−2^, and the size of the confinement area to 30 nm. A molecule that displayed at least one stabilization event, regardless of its duration, was considered stabilized.

### Monte Carlo simulations of diffusion and capture of channels

The simulation script was coded in MATLAB (github.com/mlrennerfr/DiffuTrapKv) and run on a personal computer. Trajectories were simulated in a 2D space (square of 15 × 15 µm) introducing rebound conditions on each side to keep all the molecules (200 in total) in the area and reach equilibrium.

Trajectories were simulated as in Kokolaki et al. (2020), with some modifications. Molecules were simulated as circles of 10 nm in diameter and could carry one or four scaffolding domains, representing two subpopulations of molecules with one or four PRC. To simulate a mixed population of channels, molecules with 1 or 4 interaction sites were simulated together with different proportions of each group (Fig. 4A).

The x and y components of the i-th displacement step in the trajectory were randomly selected from two independent normal distributions with the mean of zero and the variance equal to 2 *D*_*sim*_ *Δt. D*_*sim*_ was 0.01 µm^2^/s. Two different time steps *Δt* were used: *Δt* = 1 ms was used when no obstacle (other molecule) was present in the surroundings, and *Δt* = 0.1 ms in the vicinity of an obstacle. When one molecule hit another (mobile or immobile), it bounced back a distance equivalent to its diameter. At each time point, each binding domain of the molecule could engage a scaffolding interaction that confined the molecule in an area of 20 nm in diameter. The interaction started if the probability of interaction *P*_*bind*_ of at least one PRC domain was above a number *R* randomly generated from a uniform distribution. The probability of binding of each scaffolding domain was independent of the others. Stabilization lasted until the probability for detachment, *P*_*free*_, exceeded another random number. The molecule switched to free diffusion only if none of its PRC domain was interacting. *P*_*free*_ and *P*_*bind*_, were chosen to provide trajectories with a diffusive behavior similar to experimental ones. 3 independent simulation rounds were run for each case. The random generator was seeded with the current time to produce a different sequence of numbers each time. To obtain trajectories with similar characteristics to the experimental ones, simulated trajectories were converted to the temporal and spatial resolutions of trajectories obtained previously with SPT in the laboratory (acquisition frequency of 18 Hz, localization precision of 15 nm). The effect of the limited localization precision was simulated by adding Gaussian noise, with mean zero and variance equal to the localization precision).

### Electrophysiological recordings

Whole-cell patch-clamp recordings were made from 21-25 DIV primary hippocampal neurons maintained at 33°C in a recording medium containing (in mM): 120 NaCl, 20 D-glucose, 10 HEPES, 3 MgCl_2_, 2 KCl, 2 CaCl_2_ at pH 7.4, complemented with synaptic blockers (in µM): 10 Na-NBQX, 50 D-APV and 20 bicuculline methochloride. Recordings were performed using borosilicate glass pipettes of resistance 4-5 MΩ when filled with an internal solution composed of (in mM): 120 K-Gluconate, 10 KCl, 10 HEPES, 0.1 EGTA, 4 MgATP, 0.4 Na_3_GTP (pH adjusted to 7.4 with KOH). Signals were acquired with a Multiclamp 700B amplifier (Molecular Devices, USA), low-pass filtered at 10 kHz, and digitized at 20 kHz. Input resistance (Rin) was determined in voltage-clamp mode by measuring the steady-state current induced by a 1 s voltage step of −5 mV. Cells were then recorded in current-clamp mode with their membrane potential maintained around −65 mV. All voltages were corrected offline for liquid junction potential ( ≈14.5 mV in our recording conditions). In case of guangxitoxin-1^E^ (GxTx) application, cells were first recorded in control conditions, then GxTx aliquots were promptly thawed before use and pre-dissolved in ACSF before bath application.

Cells displaying non-adapting, high frequency or stuttering firing patterns were assumed to be inhibitory interneurons and discarded. When applicable, Homer-DsRed fluorescence was also used to identify pyramidal neurons thanks to the presence of dendritic spines. The membrane time constant (τ) was determined in current-clamp mode by adjusting an exponential function to the capacitive response elicited by current injection (−25 pA).

Input-output curves were generated by plotting the firing frequency in response to incremental current injections of 800 ms, 25 pA steps ranging from −25 to +475 pA. AP properties were determined from the first AP elicited at rheobase. Phase plots were constructed by plotting the time derivative of the somatic membrane potential (dV/dt) versus the somatic membrane potential of the first action potential. Analyses were performed offline using home-made codes in Matlab (Goutierre et al., 2019). employing built-in functions and FMAToolbox (https://fmatoolbox.sourceforge.net/). A MATLAB App is available at github.com/mlrennerfr/AnalyseSweeps. The AHP was calculated as the subtraction between the maximum of the AP and the minimum voltage value just after. The sAHP was calculated after the last AP, as the difference between the baseline and the minimum of membrane voltage reached after the end of the injected current step. The attenuation was calculated as the difference between the first and third inter-spike intervals.

### Statistical analyses and figure preparation

Statistical analyses were done using GraphPad Prism 9 (Dotmatics, USA). The tests that were applied are detailed in the main text. Images were prepared using Gimp (https://www.gimp.org/) or Inkscape (https://inkscape.org).

## Results

Kv2.1 channels are expressed throughout the brain, particularly in cortical and CA1 pyramidal neurons (Vacher et al., 2008). Therefore, we chose cultured hippocampal neurons as our experimental model. To facilitate identification of mutant channels, we engineered fluorescently tagged chimeras of wild-type (WT) and mutated α-subunits based on a GFP-tagged channel described in the literature (O’Connell et al., 2005) that does not impact channel activity (O’Connell et al., 2010). All of our constructs contained a fluorescent protein (GFP or Dendra2) inserted into the extracellular loop between the first and second transmembrane domains. Our objective was to model the heterozygous context of patients carrying *de novo* mutations by expressing both mutant and non-mutant subunits in the same cell. Thus, neurons were co-transfected with fluorescently tagged WT or mutant forms of Kv2.1 without knocking down the endogenous WT channel. Additionally, cells were co-transfected with Homer-DsRed (Bats et al., 2007) to detect dendritic spines and consequently identify pyramidal neurons and localize dendrites.

### Y529* and R579* mutations shift Kv2.1 distribution towards dendrites and reduced the size of Kv2.1 clusters

First, we assessed the consequences of expressing mutated forms of Kv2.1 derived from patients on the subcellular distribution of the channels. Our focus was on mutations that produced subunits that were either completely (Y529* in rat sequence, corresponding to Y533* mutation in humans) or partially (R579* in rat sequence, corresponding to R583* mutation in humans) missing the PRC domain required for interaction with VAP at ER-PM junctions (Fig. 1A).

**Figure 1:**
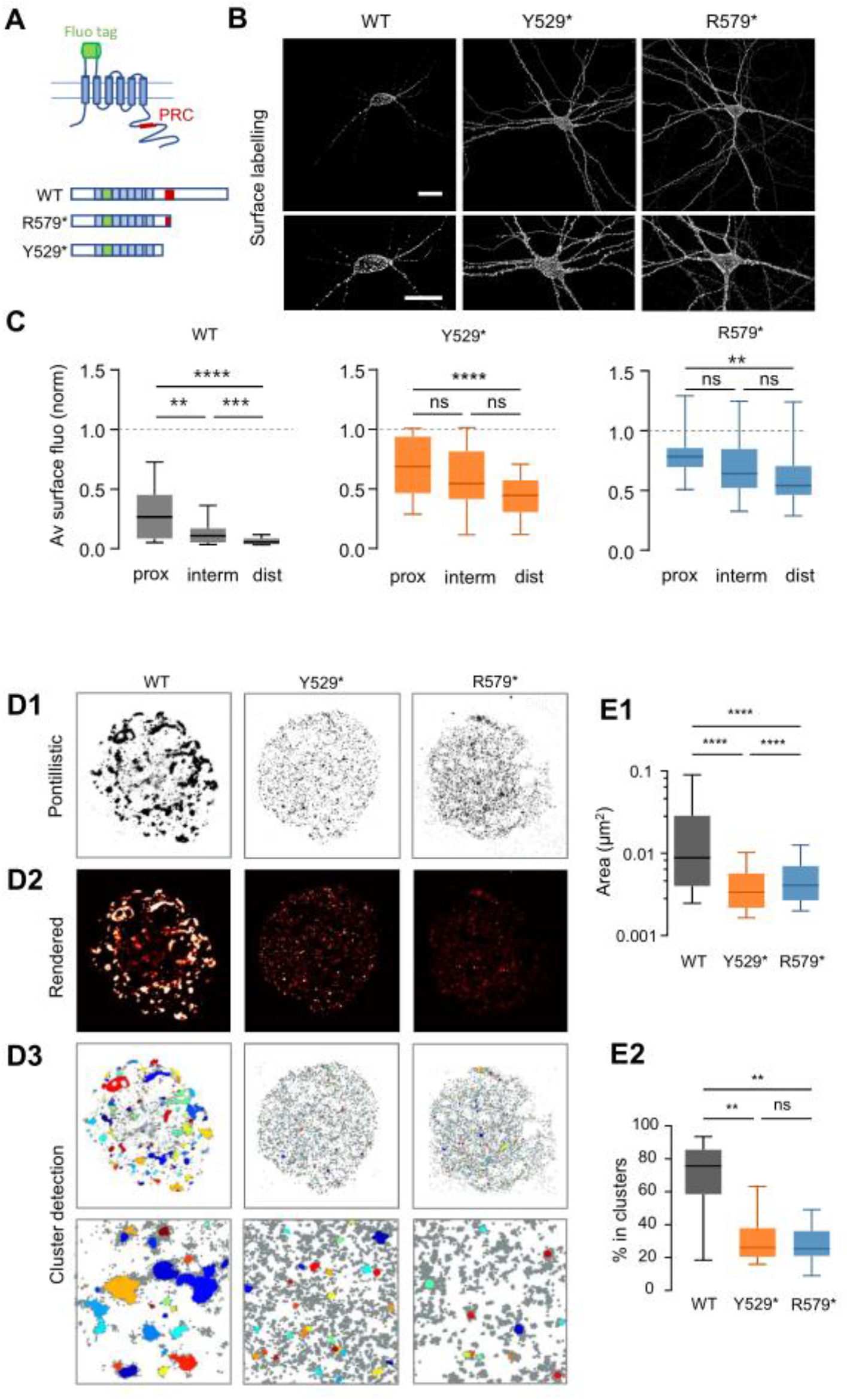
Redistribution of Y529* and R579* mutant Kv2.1 into dendritic compartments. **(A)** Schematic representation of Kv2.1 subunit chimeras (WT and mutant) indicating the locations of the fluorescent tag (GFP or Dendra2) and the PRC domain. **(B)** Top panels: confocal maximum projection images of WT and mutant Kv2.1-GFP expressed in cultured hippocampal neurons, labeled on the surface with AlexaFluor647-coupled anti-GFP nanobodies. Bottom panels: higher magnification of the regions indicated by white squares. Scale bars: 30 µm. **(C)** Average fluorescence intensity of Kv2.1-GFP staining as in A in proximal (prox., 0-50 µm from soma), intermediate (interm, 50-100 µm) and distal (dist, 100-150 µm) dendrites, normalized to the average intensity on the soma (horizontal discontinuous line). Median and 10%-90% IQR. ANOVA with Friedman tests. ns: not significant; \*\**p*<0.01; \*\*\**p*<0.001; \*\*\*\**p*<0.0001. n = 10 to 11 pyramidal neurons from 4 independent cultures. **(D)** Single-molecule localization microscopy analysis of clustering. **D1-2**: Pointillistic (D1) and rendered (D2) images of STORM detections of surface Kv2.1 GFP labeled with AlexaFluor647-coupled anti-GFP nanobodies. The focal plane corresponds to the top of the soma of cultured hippocampal neurons expressing WT and mutant channels. **D3**: Detection of clusters. The detected clusters appear in different colors. The area indicated by the black squares is shown at higher magnification in the bottom panels. Scale bars: 1 µm. **(E)** Quantification of Kv2.1 cluster’s area (E1) and percentage of detections in clusters (E2). Median and 10%-90% IQR. Kruskal-Wallis with Dunn tests. ns: not significant; \*\**p*<0.01; \*\*\*\**p*<0.0001. n = 7 to 10 pyramidal neurons from 2 independent cultures.

We verified the expression of Kv2.1 chimeras at the plasma membrane by surface staining with AlexaFluor647-coupled anti-GFP nanobodies. Confocal microscopy confirmed that the chimeric proteins were well expressed on the cell surface and that the mutated forms were distributed differently than the WT protein (Fig. 1B). Kv2.1 WT-GFP was predominantly localized in clusters at the soma and proximal dendrites. Its average surface fluorescence decreased sharply along the dendrites (Fig. 1C). In contrast, channels bearing mutant subunits exhibited a more uniform distribution between the soma and dendrites (Fig. 1C). At distal dendrites (>100 µm from the soma), the remaining average fluorescence was ~7% for WT, ~44% for Y529*, and ~63% for R579*.

Since both mutations affect the PRC domain, impaired clustering was expected. We used STORM super-resolution microscopy to analyze Kv2.1 clustering with a resolution below the diffraction limit. To ensure optimal cluster analysis conditions, we focused only on the top surface of the soma because this region has a relatively flat membrane (Fig. 1D).

Kv2.1WT-GFP formed large, micron-sized, clusters surrounded by numerous smaller ones. Overall, ~75% of detections were clustered (Fig. 1E1-2). The clustering behavior of Kv2.1 was altered in cells expressing the mutant, but not completely abolished; ~25% of the mutated channels were found in small clusters (Fig. 1D3, 1E1-2).

An important issue is whether expressing mutated Kv2.1 affects the expression and distribution of the endogenous protein. Using an antibody directed to a C-terminal epitope absent in the mutated forms, we selectively immunolabeled the endogenous Kv2.1 protein and analyzed its clustering using confocal microscopy. We found that the distribution of endogenous Kv2.1 was altered by the presence of mutants (Fig. 2A–B). While the average fluorescence of endogenous Kv2.1 decreased by ~80% from proximal to distal dendrites in non-transfected cells (Fig. 2C), the decrease was limited to ~30% in the presence of mutant channels (Fig. 2E), similar to what was observed for mutant forms (Fig. 1B).

**Figure 2.**
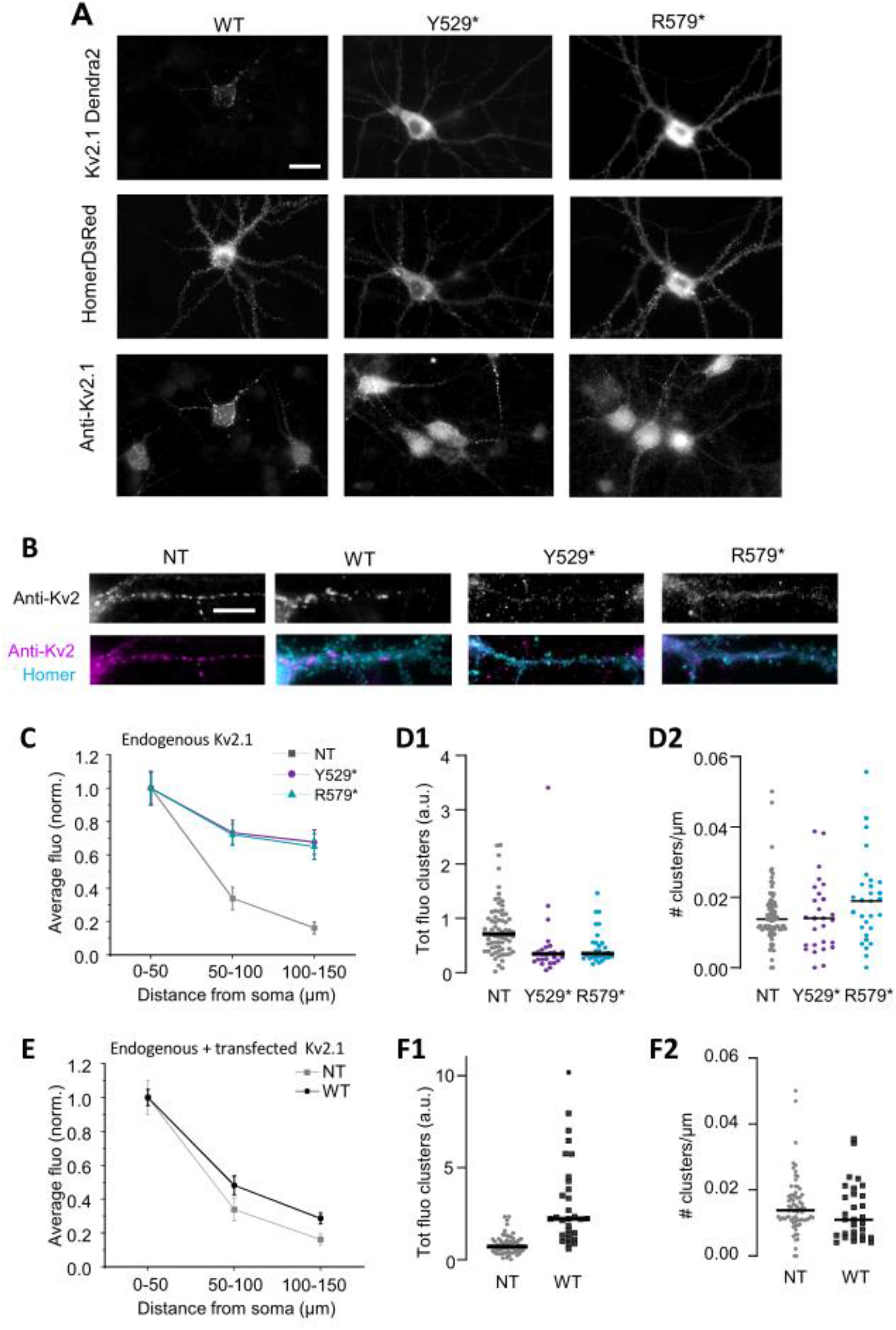
Endogenous Kv2.1 redistribution in the presence of mutated α subunits Y529* and 583*. **(A)** Epifluorescence images of hippocampal cultured neurons co-transfected with WT or mutant Kv2.1-Dendra2 (top panels) and Homer-DsRed (middle panels), immunolabeled with an anti-Kv2.1 antibody targeting the C-ter absent in mutant subunits (bottom panels). Scale bar: 20 µm. **(B)** Top panels: epifluorescence images of dendrites labelled with anti-Kv2.1 antibody (top panels) in non-transfected and transfected cells as in A. Bottom panels: merge images of anti-Kv2.1 immunolabelling (magenta) and Homer-DsRed fluorescence (cyan). Scale bar: 5 µm. **(C)** Quantification of endogenous Kv2.1 (anti-Kv2.1 staining) in non-transfected or mutant-expressing neurons: average fluorescence along dendrites (mean with SEM). **(D)** Total fluorescence of clusters (D1) and density of clusters (D2, number of clusters per unit length along dendrites). Each symbol represents a cell (horizontal line: median value). Kruskal-Wallis with Dunn tests; ns: not significant; \**p*<0.05; \*\*\**p*<0.001. **(E-F)** Quantification of anti-Kv2.1 staining (endogenous and transfected) in non-transfected or Kv2.1WT expressing neurons: average fluorescence along dendrites (E, mean with SEM), total fluorescence of clusters (F1) and density of clusters (F2, number of clusters per distance unit along dendrites). In F1 and F2 each symbol represents a cell (horizontal line: median value). **(D-F)** n = 27 to 30 pyramidal neurons from 3 independent cultures. Mann-Whitney (E) or Kruskal-Wallis with Dunn tests (F1,2). ns: not significant; \*\*\*\**p*<0.0001.

The clustering of endogenous channels was also reduced in the presence of mutated forms (Fig. 2D1). There were no significant changes in cluster density; therefore, mutations reduced cluster size without splitting them (Fig. 2D2). Thus, expression of mutant subunits exerted a dominant-negative effect on the distribution of endogenous Kv2.1, likely through the formation of heterotetramers of endogenous and mutated Kv2.1 subunits.

It has been reported that overexpressing wild-type (WT) Kv2.1 channels increases clustering (Antonucci et al., 2001). We confirmed this result by observing larger clusters in cells expressing WT Kv2.1-GFP with no significant changes in distribution along the dendrite (Fig. 2E-F). Taken together, these results demonstrate that expressing α subunits carrying or lacking the PRC domain can shift the equilibrium between clustered and non-clustered channels.

### Kv2.1 channels are immobilized both in and out of clusters, and Y529* and R579* mutations decrease their stabilization

The fluid nature of biological membranes at physiological temperatures suggests that clusters of membrane molecules form through protein-protein interactions that immobilize or stabilize the molecules via a diffusion-capture mechanism (Ribrault et al., 2011; Renner et al., 2017). Clustered Kv2.1 interacts with VAP at plasma membrane-ER junctions through a sequence in their PRC domain (Kirmiz et al., 2018). Thus, VAP act as a scaffold that stabilizes and clusters channels. Analyzing Kv2.1 lateral diffusion and detecting transient immobilization events (stabilizations) provides insight into this interaction (Renner et al., 2017). We used quantum dot-based single particle tracking to analyze the movement of WT or mutated Kv2.1-GFP molecules at the plasma membrane. The Kv2.1-GFP molecules were labeled with quantum dots (QDs) coupled to an anti-GFP antibody (QD-GFPGPI; see Fig. 3A). The QD-GFPGPI were detected with a localization accuracy of 20–30 nm and tracked at 18 Hz. As a negative control, we analyzed the diffusion of a non-related chimeric molecule, a1m (an engineered, nonfunctional version of the glycine receptor lacking intracellular domains), which does not display any scaffolding interactions (Meier et al., 2001). Thus, it reveals constraints to diffusion independent of specific stabilizations.

**Figure 3:**
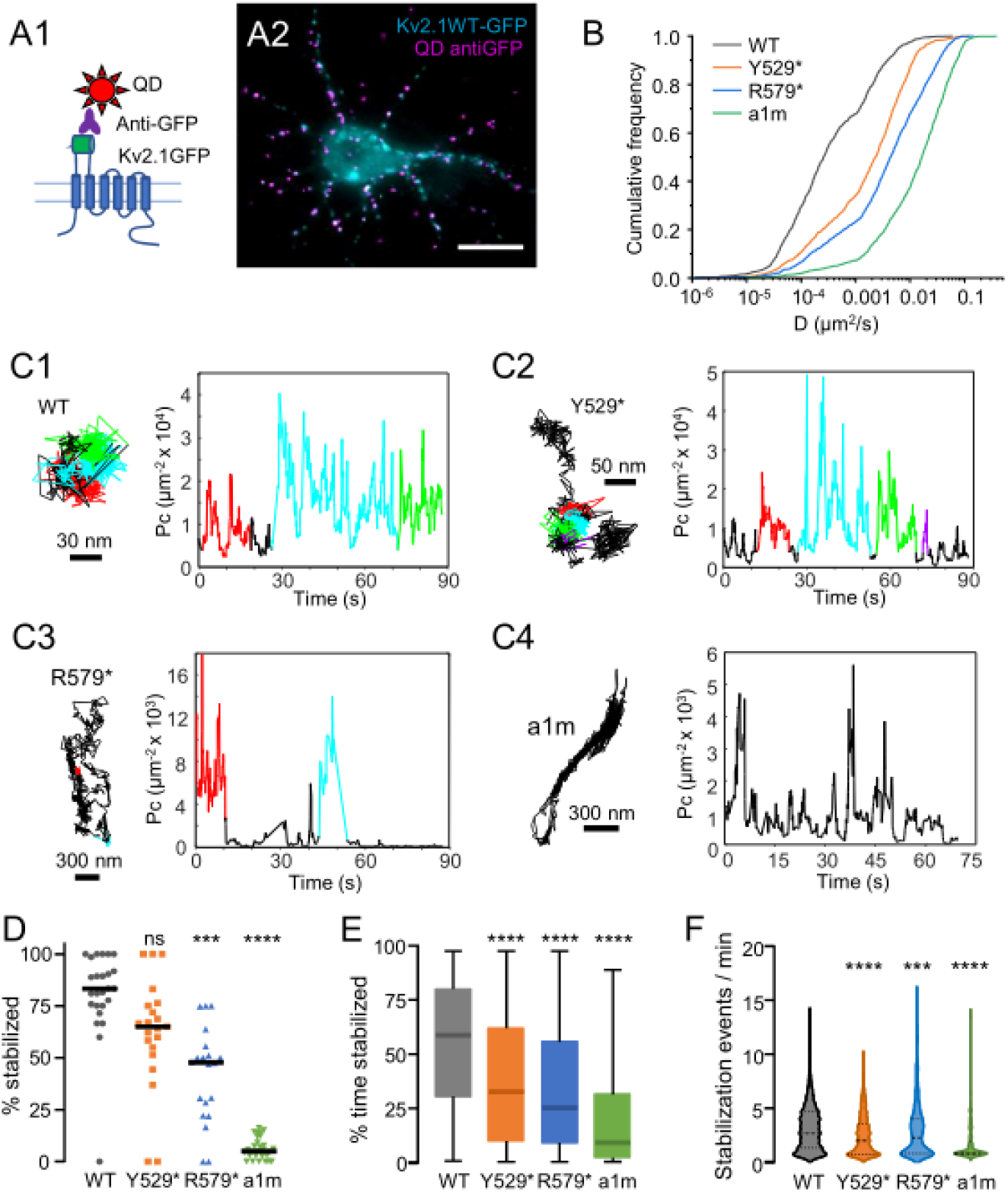
Membrane Kv2.1 channels carrying mutated subunits display higher lateral diffusion. **(A1)** Schematic representation of quantum dots labelling of Kv2.1. **(A2)** Epifluorescence image of Kv2.1WT-GFP expressing neuron (cyan) labelled with QD-coupled anti-GFP antibodies (magenta). Scale bar: 10 µm. **(B)** Cumulative frequency distributions of the diffusion coefficient (*D)* of Kv2WT-GFP (black), Kv2Y529*-GFP (red), Kv2R579*-GFP (blue), and a1m (green) trajectories. Kruskal-Wallis with Dunn tests. *p*<0.0001, n = 974-1658 trajectories, 3 independent experiments. **(C)** Examples of trajectory reconstruction with detection of stabilization events by *Pc* analysis. Representative trajectories of QD-bound Kv2.1WT (C1), Kv2.1Y529* (C2) Kv2.1R579* (C3) or a1m (C4). In color: segments recognized as stabilization events (one color per event) following *Pc* analysis. Left: position of the stabilization events on the trajectories. Right: *Pc* metric in time for the trajectories on the left, showing the detection of stabilization events based on the thresholding of *Pc*. Note the different scale of *Pc* values between the examples. **(D)** Percentage of trajectories with at least one stabilization event (independently of the duration of the event). Each symbol represents one recording. Horizontal lines: median value. Kruskal-Wallis with Dunn tests. ns: not significant; *p<0.05; ***p<0.001; ****p<0.0001, n = 20-30 recordings. **(E)** Percentage of time spent in stabilization events (median and 10%-90% IQR). **(F)** Number of stabilization events per minute (violin plots). Kruskal-Wallis with Dunn tests. ns: not significant; *p<0.05; **p<0.01; ***p<0.001; ****p<0.0001.

**Figure 4:**
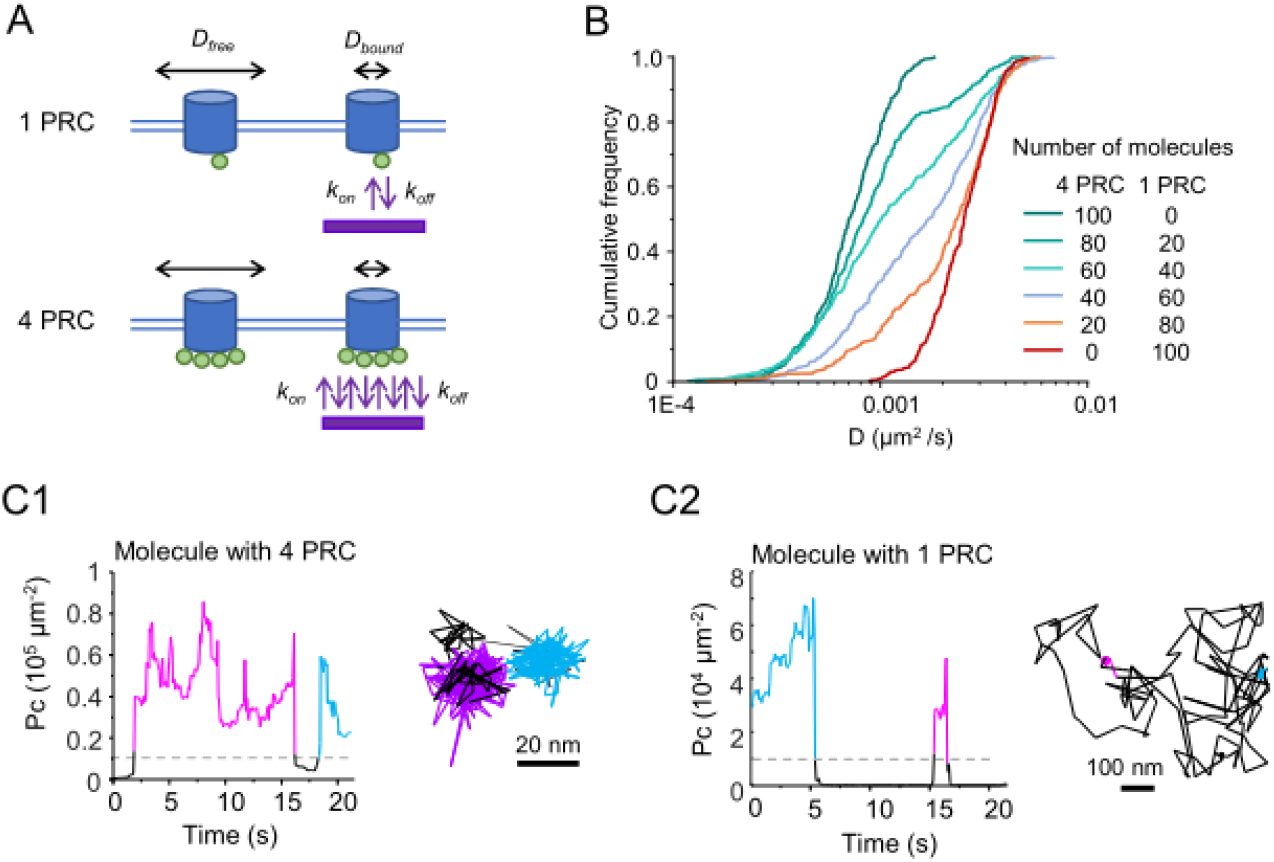
Diffusion and Pc analysis on simulated trajectories. **(A)** The diffusion and binding to immobile interactors were simulated for molecules bearing 1 or 4 binding sites (PRC), in different proportions. **(B)** Cumulative frequency distributions of the diffusion coefficient (*D)* of 6 different combinations of the aforementioned molecules. **(C)** Examples of simulated trajectories of a molecule with 4 PRC (C1) or 1 PRC (C2) with the corresponding *Pc* analysis. The immobilization periods (bound state) appear in color. In black: periods of free diffusion.

The distribution of the diffusion coefficient (*D*) of the mutated channels showed a median value that was approximately one order of magnitude higher than that of the Kv2.1WT-GFP (Fig. 3B), which is consistent with the absence of the VAP-interacting sequence. Interestingly, the shape of the curves indicated the presence of two subpopulations: one immobile or slowly moving (*D* <0.001 *µ*m^2^/s) and one more mobile. The proportion of molecules belonging to each group varied among the molecules analyzed: 62.85% ± 12.25%, 33.34% ± 9.28%, and 19.68% ± 5.87% (mean ± SD) for Kv2.1WT-GFP, Kv2.1Y529*-GFP, and Kv2.1R579*-GFP, respectively; and 6.89% ± 2.42% for a1m.

We used a method of analysis developed in the lab to detect and quantify stabilization events (Renner et al., 2017). The packing coefficient (*Pc*) parameter provides an estimate of the degree of free movement a molecule exhibits over time, independent of its global diffusivity. In the case of transient immobilization, *Pc* increases temporarily. Therefore, setting a threshold on *Pc* can detect and localize temporary stabilization events in both time and space (Fig. 3C1-4; Renner et al., 2017). Both WT and mutant subunits displayed stabilization events, but these events were often separated by free diffusion periods only in the case of mutant subunits.

We defined a molecule as “stabilized” if it displayed at least one stabilization event during the recording. Most WT channels were stabilized (~83%, see Fig. 3D) and remained stabilized for ~55% of the recording time (median value, see Fig. 3E). Consistent with the observed distribution of *D*, a smaller proportion of mutant channels were stabilized (~65% for Kv2.1Y529*-GFP and ~47% for Kv2.1R579*-GFP). However, both still displayed significantly higher stabilization levels than the negative control, a1m (~5%). The duration of each event varied greatly and did not differ significantly among groups (not shown). Most WT channels displayed more frequent stabilization events than mutated channels (Fig. 3F). Nevertheless, channels bearing mutant forms could still be stabilized. However, they were less likely to be trapped by a scaffolding interaction.

These results could be explained by the variable number of PRC domains per molecule. We hypothesized that if scaffolding interactions were labile, PRC domains would frequently detach from VAP, thereby freeing the channel. However, having several PRCs per molecule would overcome this situation by increasing the probability that at least one PRC would interact at any given time. To test this, we simulated the diffusion and stabilization of Kv2.1 with 1 or 4 interaction domains per molecule (Fig. 4A), then mixed these groups at different proportions (Fig. 4B). As expected, the population consisting of molecules with four interaction domains displayed the lowest *D* values and was often stabilized (Fig. 4B and C1), whereas the diffusion of a population of molecules with one domain was much faster and was rarely immobilized (Fig. 4B and C2). As with the experimental data, the distribution of D in the mixed groups showed two groups with different mobilities, depending on the relative amount of each group (Fig. 4B).

Overall, these results suggest that the scaffolding interactions that stabilize Kv2.1 were labile, but the presence of multiple PRCs in the tetramer ensured long-lasting stabilization of the channel.

Since ~80% of wild-type (WT) channels reside in clusters, we expected a small subpopulation of mobile molecules outside of clusters. We sorted trajectories into those belonging to clusters (defined by GFP fluorescence) and those not (Fig. 5A). As expected, the distribution of *D* for trajectories outside of clusters shifted to the right (Fig. 5B), and there was a significant reduction in stabilizations outside of clusters. However, Kv2.1 WT-GFP was still mostly stabilized without colocalizing with observable GFP clusters (Fig. 5C1), suggesting that Kv2.1 establishes scaffolding interactions outside of clusters that are detectable by wide-field fluorescence microscopy.

**Figure 5.**
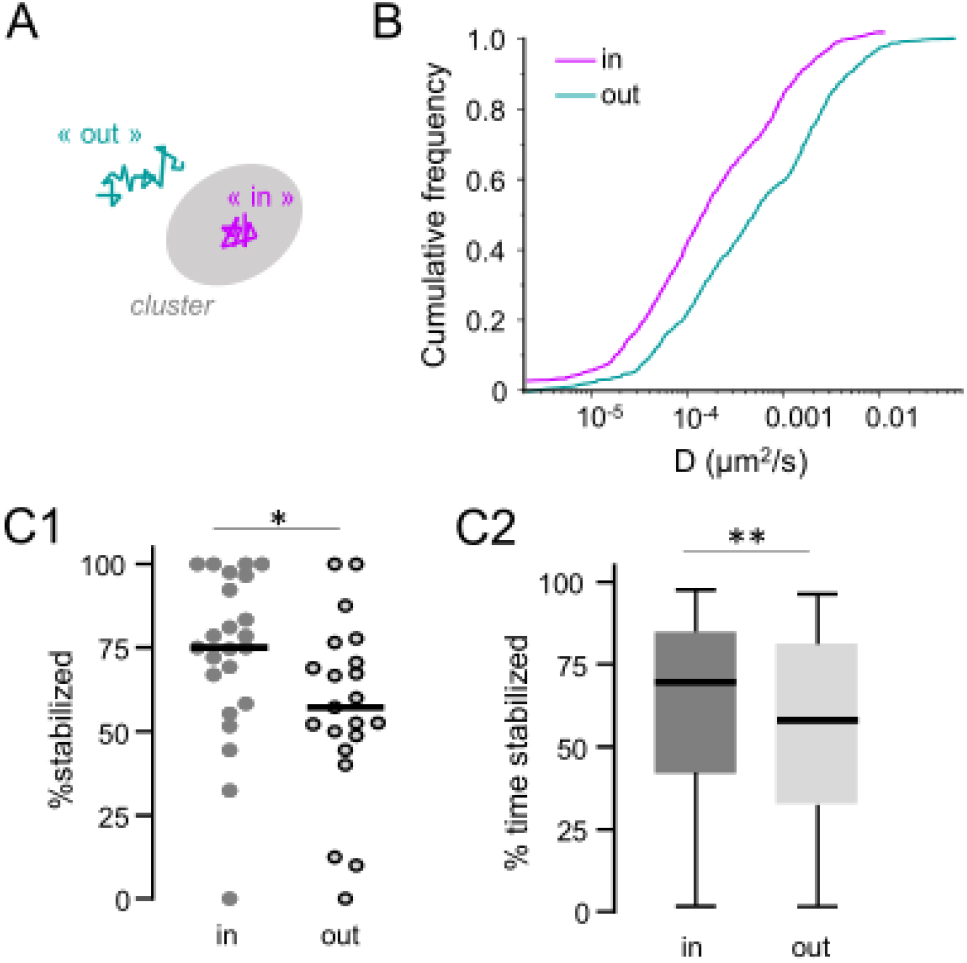
Lateral diffusion of Kv2.1WT-GFP in and out of clusters. **(A)** Schematic representation of ‘in’ and ‘out’ groups of trajectories. **(B)** Cumulative frequency distributions of (*D)* for Kv2.1WT colocalizing (in) or not (out) with Kv2.1WT-GFP fluorescent clusters. Mann Whitney test. *p*<0.0001, n = 747-1095 trajectories. **(C)** Analysis of stabilization. Mann-Whitney test. \**p*<0.05, \*\**p*<0.01, n = 23 recordings. **(C1)** Percentage of stabilized trajectories for the data in panel B. Each symbol represents one recording. **(C2)**. Percentage of time spent in stabilization events for the trajectories in B (median and 10%-90% IQR). Horizontal line: median. Data are from three independent neuronal cultures.

### Neuronal excitability is affected by Kv2.1 overexpression but not clustering

Since the conductance of Kv2.1 channels is affected by their surface density (Fox et al., 2013), we hypothesized that mutant, non-clustered channels could decrease cell excitability. We performed whole-cell patch-clamp experiments on both transfected and non-transfected neurons while synaptic transmission was pharmacologically blocked. Most of the recorded cells were identified as “regular-spiking” neurons. Only cells exhibiting the characteristics of pyramidal neurons were included in the analysis (including transfected cells with dendritic spines, as detected by Homer-DsRed puncta).

The passive properties of the recorded cells fell within the typical range described for pyramidal neurons. Input resistance ranged from 227 to 566 MΩ, the membrane time constant ranged from 26 to 77 ms, and the membrane capacitance ranged from 85 to 154 pF (25th to 75th percentile, n = 30). Cells expressing WT or mutant channels had passive properties similar to non-transfected cells (data not shown). Only cells that responded to prolonged, suprathreshold current pulses of increasing amplitude were included in the analysis. These cells displayed trains of action potentials (AP) of increasing frequency when stimulated by increasing steps of depolarizing current (Fig. 6A). No difference was observed in the characteristics of the first AP among the groups (Fig. 6B-C). The excitability of cells expressing mutant channels was not significantly different from that of cells expressing WT Kv2.1 (Fig. 6D). Our results show that changes in the distribution of Kv2.1 did not translate into changes in intrinsic excitability.

**Figure 6.**
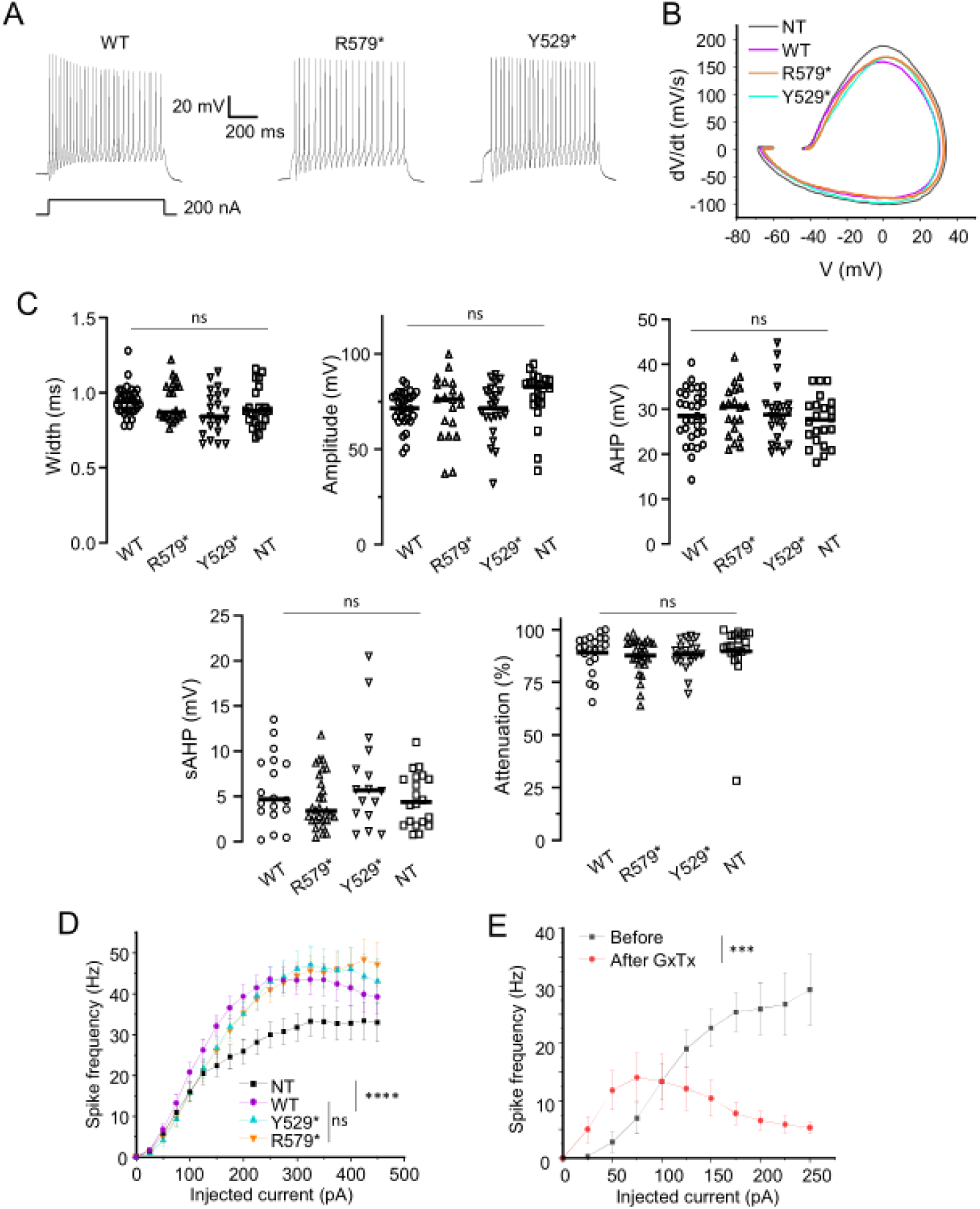
Neuronal excitability is increased by Kv2.1 WT or mutant overexpression. **(A)** Representative membrane voltage traces in transfected pyramidal neurons as indicated, obtained by applying a depolarizing current pulse of 200 pA. **(B)** Mean phase plots of the first AP for the indicated groups. **(C)** Characteristics of the first AP: width, amplitude, AHP, sAHP and attenuation, for the indicated groups. Kruskal-Wallis test, ns: not significant. **(D)** F-I curves for the indicated groups. ANOVA with Friedman test, ns: not significant, ****: p<0.0001) **(E)** F-I curves for non-transfected neurons before and after GxTx application. Wilcoxon matched pairs, ***: p<0.001. **B-D:** n = 20-31 neurons from 4 independent cultures. **E:** n=9 neurons from 2 independent cultures.

Unexpectedly, neurons expressing both WT and mutant Kv2.1 were more excitable than non-transfected cells (Fig. 6D). We checked whether the increase in excitability was due to a reduction in Kv2.1 currents in transfected cells. We analyze the excitability of the same neurons before and after application of Kv2 specific blocker Guanxitoxin-1E (GxTx, Liu & Bean 2014). GxTx had a completely different effect. It initially increased AP frequency, but cells rapidly entered depolarization block (Fig. 6E). Therefore, blocking Kv2 channels reduced the maximum firing frequency of pyramidal cells. These results suggest that transfected cells reached higher firing frequencies likely due to the overexpression of Kv2.1 due to transfection.

## Discussion

Kv2 channels, particularly Kv2.1, are widely distributed in the brain and are a major component of neuronal delayed rectifier K^+^ currents. Despite their prominence, the physiological roles of Kv2 channels are less well understood than those of other voltage-gated K^+^ channels. This is probably due to the relatively recent characterization of their specific blocker, GxTx (Hönigsperger et al., 2017; Liu & Bean, 2014).

The identification of Kv2.1 mutations in patients with epileptic encephalopathies has sparked new interest in these channels. Kv2.1 is unique among voltage-gated potassium channels due to its localization in clusters. We characterized two *de novo* mutations that partially (R579*) or completely (Y529*) eliminate the PRC domain, which is responsible for clustering. Patients expressing these mutations exhibit mild epileptic phenotypes accompanied by cognitive impairments and autistic behavior (Bar et al., 2020).

Our results confirm the importance of the PRC domain for Kv2.1 subcellular distribution and clustering. Despite the lack of PRC domains, channels carrying mutated subunits still showed residual clustering, most likely due to heterotetramerization with endogenous subunits. This is likely the situation found in patients because *de novo* mutations affect only one allele of the *KCNB1* gene. Importantly, endogenous Kv2.1 distribution was altered in the presence of mutated subunits, suggesting that Y529* and R579* act as dominant-negative subunits for Kv2.1 localization. The large majority of mutated channels were not clustered (~75%) and their distribution shifted towards dendrites. This result confirms that there is a signal sequence in the portion of the C-ter lost in mutant subunits that confines channels to the somatodendritic compartment.

We distinguished two distinct populations of channels based on the distribution of diffusion coefficients. We speculate that the small proportion of faster molecules within WT channels corresponds to heterotetramers, such as Kv2.1-Kv6 complexes, which carry only two PRC domains (Möller et al., 2020). Consistent with the change in clustering behavior, mutant subunits increased the proportion of faster channels.

The proportion of slow and fast channels also correlated with cluster size, supporting the idea that cluster formation relied on a diffusion-capture mechanism enhanced by several PRC domains. SPT experiments and diffusion-capture simulations suggest that heterotetramers of endogenous and mutant subunits did not contain enough PRC domains to stabilize the channels sufficiently to form large clusters. Our results also indicate that scaffolding interactions are labile; thus, cluster formation is dependent on the number of PRC domains per molecule. For WT homotetramers, the presence of four PRC domains ensures that at least one will interact, rendering these channels nearly immobile. In contrast, channels with fewer PRC domains are less stable and prone to diffuse freely. Therefore, channels with mutant subunits would be stabilized for shorter periods of time and form only short-lived nanoclusters.

Interestingly, WT channels were strongly stabilized even if they did not colocalize with large clusters. This result could be explained by the presence of abundant scaffolding interactions not only restricted to clusters visible by regular fluorescence microscopy. In fact, super-resolution microscopy showed that a large proportion of WT channels were forming small clusters. The reported structural role of Kv2.1 at ER-PM contacts was only analyzed on large clusters thus further experiments will be needed to know whether the interaction with VAPs at ER-PM contacts is also responsible for Kv2.1 immobilization in nanoclusters. If this is the case, our results suggest that these contacts would be more abundant than expected from the distribution of large Kv2.1 clusters. Alternatively, the stabilization of Kv2.1 in nanoclusters would rely on other interaction.

It has been reported that Kv2.1 conductance is modulated by the surface density of channels in heterologous cells (Fox et al., 2013). Despite significant changes in Kv2.1 clustering and distribution, cells overexpressing mutant subunits did not exhibit significantly different excitability than cells overexpressing WT subunits. The AP characteristics were unchanged, suggesting that mutations did not modify Kv2.1 current.

Importantly, we observed that both WT and mutant subunits increased cell excitability compared to non-transfected cells. We cannot attribute this result to a reduction in Kv2.1 current because blocking Kv2 currents with GxTx reduced the maximum spiking frequency and cells entered faster into a depolarization block. Conversely, our results are consistent with the known role of Kv2 channels in facilitating high-frequency firing. Therefore, increasing the level of Kv2.1 expression can be a critical and paradoxical driver of hyperexcitability.

## Acknowledgements

We thank J. R. Martens (University of Florida, USA) for the original GFP-tagged Kv2.1 construction and the IFM imaging platform for their help with the imaging experiments.

## Author contribution

A-L. Paupiah carried out imaging and electrophysiological experiments, ran neuronal simulations and analyzed data. M. Ginisty performed electrophysiological experiments and analyzed data. M. Russeau prepared primary neuronal cultures. I. Mouktine designed and prepared plasmids. S. Levi provided the equipment for single particle tracking. JC. Poncer provided materials and equipment for electrophysiology, designed and supervised electrophysiological experiments. M. Renner directed research, wrote the codes for analysis of single particle tracking and simulations and supervised the experimental work. A-L. Paupiah and M. Renner prepared the figures and wrote the paper.

## Data availability

All data reported in this paper will be shared upon request. Any additional information required to re-analyze the data reported in this paper is available from the lead contact upon request.

